# Spatial Evolution of Information Dynamics in the Primary Motor Cortex During Reach

**DOI:** 10.1101/2022.12.23.521792

**Authors:** Wei Liang, Vasileios Papadourakis, Nicholas G. Hatsopoulos

**Author notes:** Nicholas G. Hatsopoulos, **Email:**.

## Abstract

The primary motor cortex (M1) is known to be spatially organized in terms of muscles and body parts, notwithstanding some overlap. However, the classical view is largely static, neglecting potential representational changes on a fast time scale. To address potential representational dynamics, we probed the mutual information between M1 signals recorded across the cortical sheet and electromyography (EMG) activity from a set of muscles while a macaque monkey performed planar reaches. Here, we demonstrated that the spatial organization of the M1 encoding was in fact quite dynamic throughout the course of a reaching movement such that a given cortical site maximally encoded different muscles at different times. Despite these rapid representational changes, a proximal-to-distal gradient of the upper limb representation was preserved particularly close to movement onset. Representation was most stable close to movement onset, with characteristic topographic maps, possibly serving functional needs. This study bridges the important gap between flexible motor encoding and static motor maps, emphasizing the importance of considering space and time together in motor representation studies.

## Introduction

For over a hundred years, people have examined whether and how movement representations in the primary motor cortex are spatially organized (Jackson, 1867; Fritsch & Hitzig, 1870; Ferrier, 1886). Various movement relevancies have been investigated, largely assuming a largely static nature of representation over short period of time.

Early maps were generated with surface electrical stimulations, including the simiusculus plotted by Woolsey and colleagues in monkeys (Woolsey, 1952), as well as the homunculus plotted by Penfiled and colleagues in humans (Penfield & Boldrey, 1937; Penfield & Rasmussen, 1950). In those maps, an ordered sequence of body parts (foot, legs, trunk, shoulder, upper limb, hands, face and mouth) was illustrated from medial to lateral cortical sheet, giving an oversimplified impression that the mapping from sites on the primary motor cortex to body parts were clear-cut. Even though the authors did note a certain degree of overlap among body part representations, this fact has not been fully appreciated even today.

With the development of intracortical microstimulation (ICMS) (Asanuma & Sakata, 1967), researchers were able to perform more focal stimulation using penetrating microelectrodes. Park and colleagues (2001, 2004) stimulated layer 5 of primary motor cortex with short-duration low-frequency ICMS, while macaques performed reach-to-grasp tasks. They confirmed the overlapping nature of representation at single sites for the upper limb area, where proximal and distal muscles were co-represented synergistically. They also found a fine horseshoe-shaped spatial map within the forelimb representation, where shoulder and elbow areas surrounded a transitory coactivation zone, which in turn surrounded hand areas.

As people began to explore additional spatial organization, Graziano and colleagues (2002, 2005) found a systematic spatial organization of ethological actions using longer ICMS trains (500ms) where actions such as the hand moving to lower peripersonal space, hand manipulation in central peripersonal space, chewing/licking were encoded sequentially from medial to lateral portions of M1. They also found that the final arm and hand postures (mainly in the vertical and horizonal axes) were represented at distinct cortical regions. Apart from the above, no global cortical spatial organization has been found with other functional properties of movement, such as velocity, acceleration, force, or change of force (Georgopoulos et al., 2007; Stark et al., 2009; Evarts, 1968; Humphrey et al., 1970).

Even though we know that cortical representation maps are malleable in the long term due to development, learning or injury (Chakrabarty & Martin, 2000; Glees & Cole, 1950; Nudo et al., 1996), the spatial structure of movement representations during a given behavior is largely assumed to be invariant in the examples above, perhaps due to limitations of the ICMS paradigm. This static assumption seems unwarranted, especially considering that movement encoding itself is flexible and dynamic within very short time (Ben-Shaul et al., 2003, 2004; Churchland & Shenoy, 2007; Hatsopoulos et al., 2007; Rokni et al., 2007). Moreover, in the few studies that looked at space and time simultaneously in the representation, no global spatial structure on the cortical sheet has been discovered accompanying representational changes (Ben-Shaul et al., 2003; Hatsopoulos et al., 2007).

Here for the first time, we tracked the dynamics of cortical representational maps of forelimb muscles during single reaches, using mutual information to quantify the relationship between neural activity and electromyography (EMG) activity. We chose the high-gamma band signal in the local field potential as our proxy for neural activity, given its rich information and dense spatial coverage (Rickert et al., 2005). We found that cortical representation across space evolved throughout movement, but certain spatial organization principles such as proximal to distal gradient were still maintained. Those spatio-temporal representation changes were different for different movement directions, with interesting stability points, likely serving functional needs.

## Methods

### Surgery procedures

A male rhesus macaque (*Macaca Mulatta*) Bx was the subject for this study. He was implanted with bipolar electromyography (EMG) electrodes in 13 muscles from shoulder to hand (Anterior Deltoid, Posterior Deltoid, Pectoralis Major, Biceps Lateral, Biceps Medial, Triceps Long head, Triceps Lateral, Brachioradialis, Flexor Carpi Ulnaris, Extensor Carpi Ulnaris, Extensor Carpi Radialis, Extensor Digitorum Communis, Flexor Digitorum Superficialis). He was also implanted with a set of dual Utah multi-electrode arrays (Blackrock Microsystems, Salt Lake City, UT) in the left primary motor cortex, each containing 64 electrodes. Electrode length was 1.5mm, with a uniform inter-electrode distance of 4µm in the 8-by-8 grid. The arrays corresponded to the arm/hand areas of the right limb, as confirmed by electrically stimulating the cortex during surgery with surface electrodes and observing corresponding twitches prior to array implantation. All procedures regarding surgery, animal training and data collection were approved by the University of Chicago Institutional Animal Care and Use Committee (IACUC) and obey the Guide for the Care and Use of Laboratory Animals.

### Behavioral task

The animal was trained to perform a reach task in different directions, with his right upper limb constrained by a 2D exoskeletal robot (BKIN Technologies). All the reach directions were on a horizontal plane roughly at the height of his elbow when his (right) upper limb was naturally hanging. To perform the task, he moved the joystick at the tip of the robot, while receiving simultaneous visual feedback of the cursor position on the screen in front of him. The task involved a hold period for 1000ms, during which the animal held the joystick steadily, keeping the cursor within a central target. The peripheral target then appeared indicating the desired reach location. He then moved the joystick to direct the cursor to the peripheral target. Each target had a radius of 0.75cm; the distance from the central target to the peripheral target was 5.5cm. The peripheral target could appear in one of 8 possible locations on a circle (in 45° intervals clockwise from 0° to 315°, where forward movement was 0° and rightward movement was 90°). A trial was considered successful once the cursor reached the peripheral target and remained in the target for a variable period of time (400 - 600ms), upon which the animal received juice reward. Movement durations vary across reach directions, but generally ranged between 300 to 1200ms. We also had the monkey perform 2-target center-out task (where target appears in one of two directions: 45° and 225°), to get more trials out of each reach direction. This 2-direction reach task had variable hold time from 800-1200ms, but otherwise identical to the 8-direction task. There was one session for each variant of the task.

### Collection and preprocessing of neural and muscle data

EMG signals from 13 muscles were individually amplified, band-pass filtered between 0.3 Hz to 1kHz, digitalized and sampled at 2k Hz. Then for preprocessing, EMG activities were further band-pass filtered between 20 Hz to 1kHz, followed by rectification and smoothing by low-pass filtering below 10Hz. Then, envelopes were down-sampled to 200Hz. Individual trials of muscle activities were aligned on movement onset (defined as the time point where 15% of the peak speed was reached for that trial).

Neural signals were collected with Blackrock Microsystems (Salt Lake City, UT). To obtain local field potentials, neural signals from 128 channels were low-pass filtered below 500Hz with a Butterworth filter and then digitized at 2kHz. From there, high-gamma band signals were extracted by band-pass filtering within 200Hz - 400Hz with a Butterworth filter. We then computed the Hilbert transformation to obtain the amplitude envelopes. Finally the envelopes was low-pass filtered below 10Hz to match the smoothness of muscle signal, and then down-sampled to 200Hz.

### Intracortical microstimulation

Short-duration intracortical microstimulation (ICMS) was performed to map the classical somatotopic map. For each stimulation event, we stimulated an electrode for 25 biphasic pulses (cathodic first) at 333Hz. The duration of positive and negative phases were 200µs, with an interval duration of 55 µs. We started at 20µA for each electrode. If any muscle twitch was observed, we lowered the current to determine the threshold and isolate the effect; if no muscle twitch was observed, we increased the current level. If no muscle twitch was elicited at 40µA, the effect of stimulating this electrode would be considered as ‘not observed’.

### Estimation of mutual information

To comprehensively quantify the dependent relationship between two variables, we used mutual information (MutInfo). The mutual information between two continuous variables *x* and *y* is a function of the joint probability distribution and marginal distributions:

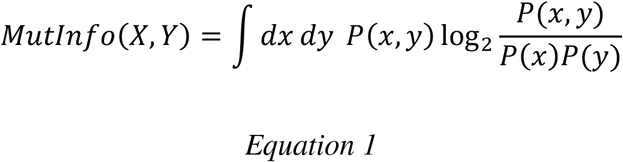

Traditional estimations used fixed or adaptive binning to extract the probabilities above (eg. Darbellay & Vajda, 1999), which are subject to non-negligible systematic errors (Kraskov et al., 2004). Instead, we used Kraskov-Stögbauer-Grassberger estimator, a binless mutual information estimator (Kraskov et al., 2004). It utilizes another formulation of mutual information, expressed by the difference between marginal and joint entropies:

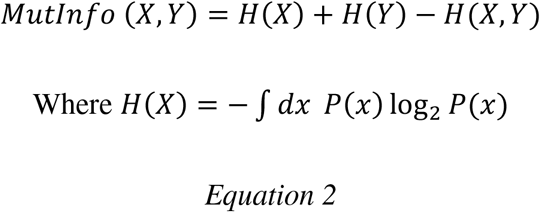

The probability distributions above were estimated based on the *k-*th nearest neighbor distances, where multivariate neighbors were defined based on the maximum norm. By varying *k*, one can adjust the spatial scales on which relationships are being probed. Important expansions of this method were made by Holmes and Nemenman (2019) which enables estimation of error bars and detection of sample-size dependent bias, facilitating the selection of *k*. Sample-size dependent bias is present when MutInfo estimations made with non-overlapping subsamples of the full data (e.g. until 10 divisions) result in much lower estimates than using the full dataset (e.g. beyond one standard error). A small *k* generally (1) enables the detection of features at a finer scale, (2) utilizes data points better and possesses less sample-size dependent bias, but (3) is also more susceptible to fluctuations in the distribution thus having a larger error bar (sometimes estimates could reach impossible negative MutInfo values within one standard error). In our case, *k* was selected to strike a balance among those considerations: we chose *k* values that resulted in acceptable standard deviations and minimal sample-size dependent bias. For all-directions pooled MutInfo computations, we used *k*=4; for direction-separated MutInfo computations, we used *k*=9. This was to deal with the increasing fluctuations in the distributions when less data points were available. Our results were qualitatively stable across a range of *k* values.

In order to track the changes of mutual information along movement execution, we divided the EMG activities into 29 sliding time windows, with 200ms window size and 50ms step size (i.e. - 400ms ∼ -200ms, -350ms ∼ -150ms, …, 1000ms ∼ 1200ms). Within each window, we consider different leads/lags of muscle and neural activity, from 600ms neural-led to 400ms muscle-led in steps of 5ms. In the direction-pooled version, mutual information was computed between neural signal and EMG for each electrode-muscle pair at each lead/lag, with all sample points of each time window from all trials; in the direction-separate version, mutual information was computed between neural signal and EMG for each electrode-muscle pair at each lead/lag, with sample points of each time window further down-sampled by 4 times (i.e. 50Hz), for trials with the same target direction. Subsampling was done to address the problem of autocorrelation in the continuous signals, especially given less diversity in trials in the case of direction-specific data. As our interests are mainly focused on relative strengths, (remaining) autocorrelation should not be a severe problem regarding mean estimates to either pooled or direction-specific variants. As for its effect on error bar estimation, we carefully mitigated it by choosing *k*s that are not too small so that our comparisons would still be meaningful even when error bars were slightly underestimated.

Key implementation code was based on Holmes and Nemenman (2019), which could be accessed at https://github.com/EmoryUniversityTheoreticalBiophysics/ContinuousMIEstimation.

### Computation of MutInfo-based somatotopic map

At each particular *k* and subsampling ratio (and for each reach direction, in the case of direction-specific mutual information), we arrived at a 4-dimensional tensor of mutual information of shape 29 (# of absolute time windows) × 13 (# of muscles) × 201 (# of relative times) × 128 (# of electrodes).We selected the range of relative time from -300ms to 300ms and then computed the maximal MutInfo over the axis of relative times, arriving at a 3-dimensional tensor of 29 (# of absolute time windows) × 13 (# of muscles) × 128 (# of electrodes).

To arrive at the full map, for each absolute time window and for each electrode, we picked the muscle that had the maximal mutual information out of the 13 muscles, and used the color corresponding to that muscle to represent that electrode on the cortical map. To arrive at the ‘motor’ map from the full map, we only kept electrodes where maximal mutual information values were obtained at negative relative times (i.e. neural activity preceding EMG). To arrive at the ‘sensory’ map from the full map, we only kept electrodes where maximal mutual information values were obtained at positive relative times (i.e. EMG preceding neural activity).

### Computation of representation evolution (non-spatial view)

Like above, at each particular *k* and subsampling ratio (and for each reach direction, in the case of direction-specific mutual information), we arrived at a 4-dimensional tensor of mutual information of shape 29 (# of absolute time windows) × 13 (# of muscles) × 201 (# of relative times) × 128 (# of electrodes).We selected relative time range from -300ms to 300ms and then computed the maximal MutInfo over the axis of relative times, arriving at a 3-dimensional tensor of 29 (# of absolute time windows) × 13 (# of muscles) × 128 (# of electrodes). We then computed the space-independent representation using two methods: summing-up or winner-take-all.

With the summing-up method, for each absolute time window and for each muscle, we summed up all the mutual information values for all electrodes, arriving at a matrix of mutual information, of shape 29 (# of absolute time windows) × 13 (# of muscles). Next, muscles were grouped according to the representation classification of interest (i.e. body parts — shoulder/elbow/wrist/hand, FE — flexion/extension, or both body part and FE — eight combos). Finally, for each absolute time window, mutual information values for muscles of the same group were added up, then divided by the number of muscles in that group. This last step of division was to ensure that representation intensity was not affected by the different number of muscles we had sampled in each group. We ended up with a representation matrix of shape 29 (# of absolute time windows) × 2/4/8 (# of groups).

With the winner-take-all method, for each absolute time window and for each muscle, we counted the number of electrodes that had this muscle as the top-represented muscle, arriving at a matrix of electrode counts, of shape 29 (# of absolute time windows) × 13 (# of muscles). Next, muscles were grouped according to the representation classification of interest like above. Finally, we adjusted the electrode count by dividing by the number of muscles in that group, arriving at representation matrices as explained above.

### Tensor decomposition

For mutual information computed separately for eight directions, we reorganized tensor into a 3-dimensional tensor of shape [# of directions × (# of muscles × # of electrodes) × (# of absolute time windows × # of relative times)]. To understand potential low-dimensional structures of the mutual information tensor, we performed a form of non-negative tensor factorization, namely CANDECOMP/PARAFAC (canonical decomposition/ parallel factors, CP) decomposition (Kim & Park, 2012; Kola et al., 2017), which approximates the tensor as a sum of outer products (i.e. factors) of 3 non-negative vectors (one for each dimension). Code is available at https://github.com/kimjingu/nonnegfac-matlab/blob/master/ncp.m and https://www.tensortoolbox.org. The performance of reconstruction is measured by the relative error, which is the ratio between the Frobenius norm of the error tensor and the Frobenius norm of the original tensor, with the error tensor being the difference of the original tensor and the reconstructed tensor. Number of factors were selected based on a balance of low relative error and interpretability.

## Results

### Somatotopic representation is dynamic during reach

During the center-out task, our monkey subject performed constrained reaches towards one of eight directions on the horizontal plane (Fig. 1a), while his EMG activity and high-gamma LFP activity were extracted. To start with, we wanted to obtain a general understanding of the evolution of representation around movement onset. For that, we computed the mutual information between each muscle and electrode pair, using data from all reach directions. For example, for the pair of Anterior Deltoid (Fig. 1b) and electrode 101 (Fig. 1c) using a -80ms lag (neural activity preceding muscle activity), mutual information changed throughout the reach course, reaching the peak at around -50ms relative to movement onset. For the same muscle with another electrode (electrode 9, Fig. 1e) also using -80ms lag, the temporal evolution of mutual evolution followed a slightly different time course, reaching peak at movement onset (Fig. 1f). Nevertheless, there was a general trend for information to be initially low, then increase transiently before returning to baseline levels. For most of them, there was a sharp increase in mutual information around movement onset.

**Figure 1.**
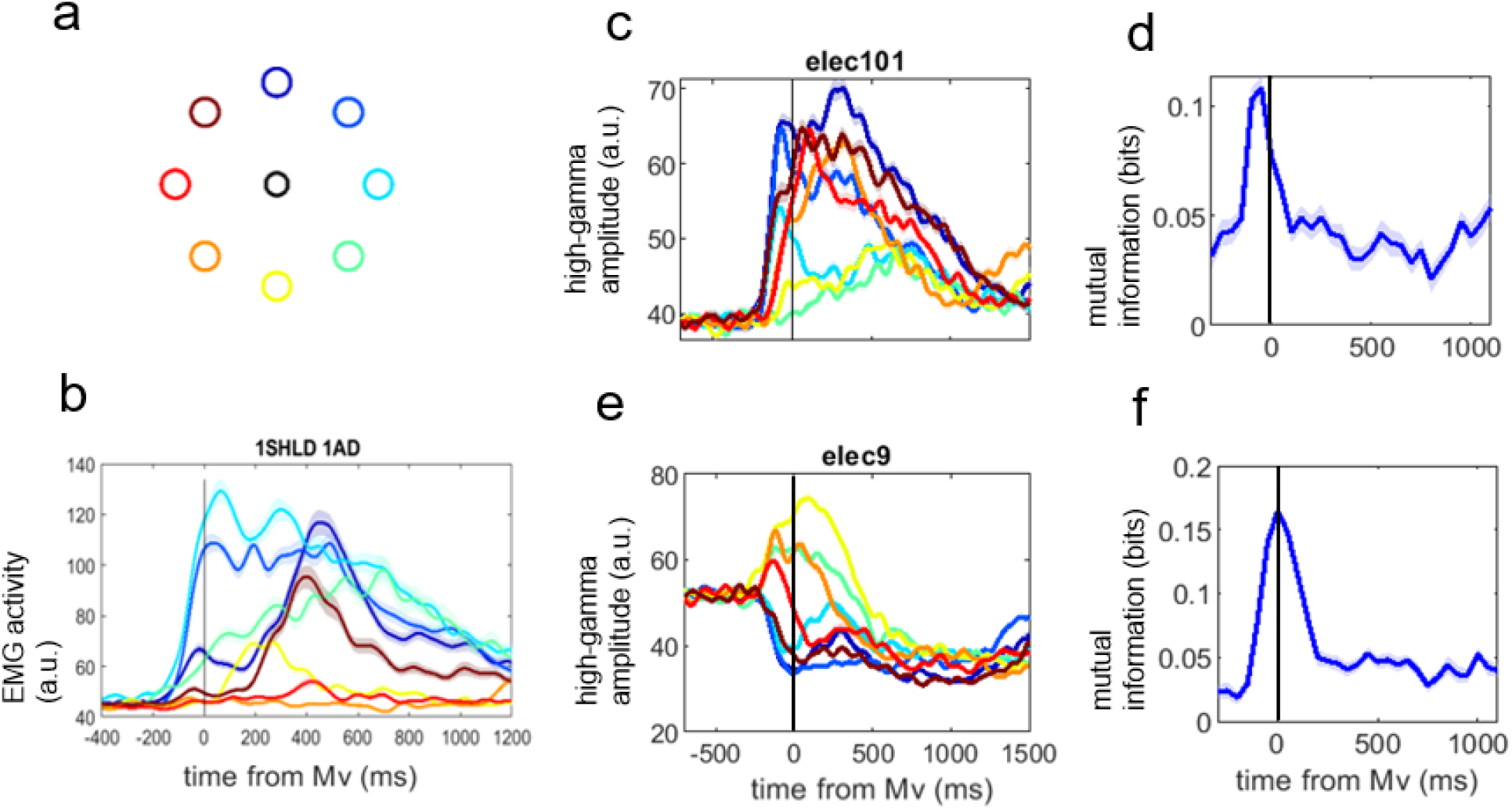
Different mutual information evolutions for different muscle-electrode pairs. **a**. movement task was a constrained reach task from the central black target to one of eight colored targets on the periphery. **b**. EMG activity of a shoulder muscle Anterior Deltoid, averaged for each reach direction separately. Color of traces were the same as target illustration in **a**. Shaded area represented standard error of mean (SEM). **c**. amplitude of high-gamma envelope at electrode 101, averaged for each reach direction separately. Color of traces were the same as target illustration in **a**. Shaded area represented SEM. **d**. mutual information computed for muscle in **b** and electrode in **c** at -80ms relative timing (neural activity preceding muscle activity by 80ms), separately for each 200ms-long time window along movement course. Trials from all directions were considered for this figure. Shaded area represented SEM. **e**. same as **c** but for another electrode (electrode 9). **f**. same as **d** but for another pair: muscle in **b** and electrode in **e**. Similar rise of mutual information was present upon movement onset but at a slightly later time than in **d**.

Next, we aimed to visualize the representation change on the cortical sheet. For each electrode, we considered the muscle that was represented most (i.e. the muscle with which this electrode had the most MutInfo) in a certain time window. There are three versions of somatotopic maps: full map (Fig. 2a), ‘motor’ map (Fig. 2b) and ‘sensory’ map (Fig. 2c). The ‘motor’ map only displayed electrodes whose maximal MutInfo were obtained when neural activity preceded muscle activity; while the ‘sensory’ map only displayed electrodes whose maximal MutInfo were obtained when neural activity lagged muscle activity. The full map combined both maps and further included a few electrodes where the maximal MutInfo were obtained with simultaneous muscle and neural activity. Note that we are not implying any direct causality here, as that is beyond the connotation of mutual information.

**Figure 2.**
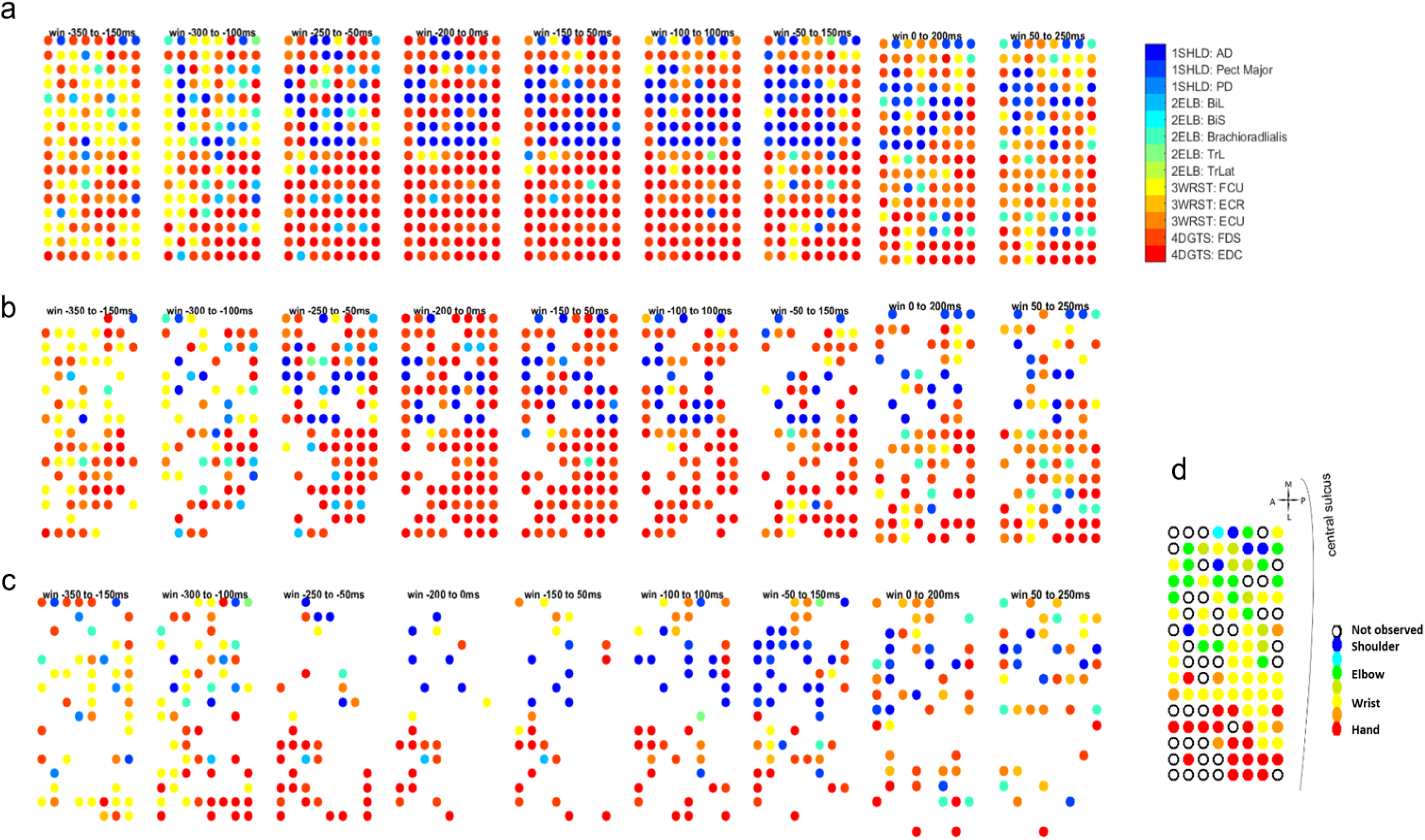
Mutual information based somatotopic maps were dynamic and different from ICMS-based map. **a**. mutual information based somatotopic map - full map. Each circle was color-coded by the muscle that was most represented at this site at this time window in the movement course. **b**. mutual information based somatotopic map – ‘motor’ map. Same as **a** but only showing electrodes that obtained the maximal mutual information at relative times where neural activity preceded muscle activity. **c**. mutual information based somatotopic map – ‘sensory’ map. Same as **a** but only showing electrodes that obtained the maximal mutual information at relative times where neural activity lagged muscle activity. **d**. ICMS-based somatotopic map, color-coded by the body part that twitched during electrical stimulation.

In all those maps, representations of the same muscle were distributed across a relatively large area of the cortical sheet, but also possessing preferred congregation area. Looking at the full map (Fig. 2a), a majority of wrist representation was present during the hold preparation period. Then closer to movement onset, a lot of those wrist representation was replaced by a separation of shoulder and hand representation, roughly following the proximal-distal gradient in the ICMS-based somatotopic map. As the movement unfolded, more wrist and elbow representation started to come in. As for the ‘motor’ and ‘sensory’ maps, during hold we observed slightly more ‘motor’ sites than ‘sensory’ sites, then closer to movement onset, there was a sharp increase of ‘motor’ sites, which gradually receded as movement unfolded, giving place to ‘sensory’ sites. Interestingly, the representation in the corresponding ‘motor’ and ‘sensory’ maps were quite representative of the full map, with similar ratios of muscle preferences and locations. This implied the integrative nature of the ‘motor’ and ‘sensory’ sites in the primary motor cortex, mostly likely from one continuity.

Note that despite the constant evolution, the MutInfo-based somatotopic maps never resembled the ICMS-based somatotopic map. In general, MutInfo-based maps had less distinct boundaries between body parts and were more distributed; the ratio of body part representation constantly changed, and a certain electrode could represent different muscles at different times.

### Representation evolution is movement-specific and widely distributed

Beyond general representation change throughout movement obtained using a pool of heterogenous reaches, we wanted to know whether different movement directions possessed different representational dynamics. For that, we computed mutual information for each muscle-electrode pair at each relative times at each absolute time windows, separately for each reach direction.

Before diving into the many new sets of somatotopic maps, for simplicity we first illustrated the changes of representation strength in a space-independent way. Here, we added up mutual information obtained from different electrodes for the same muscle, then further grouped the muscles by their body locations and flexion/extension (i.e. polarity) function. (Note that this summation operation on mutual information is not perfect — see Discussion for further considerations and justifications). The summed mutual information was then adjusted according to the number of muscles in that group, to reduce the effect of biased muscle sampling (see Methods for details). Fig. 3 shows the resultant dynamic changes of representation strength through reaches of different directions. Adjacent directions tended to have similar dynamical patterns. Most muscle categories obtained peak representation strength around movement onset for most directions, with some exceptions: shoulder flexors and wrist extensors peaked later for forward and leftward directions (Fig. 3 a, g, h); a different type of representation strength evolution was present for inward and inward-rightward directions (Fig. 3 d and e), characterized by slow rise and delayed (multiple) peaks. While digit flexors were the most represented group closer to movement onset for most directions, shoulder extensors were most represented for inward and leftward directions (Fig. 3 e-g). Notably, there was great consistency between sessions, as results obtained from a 2-direction reach session on another day showed very similar evolution of representation strengths (insets in Fig. 3b and f).

**Figure 3.**
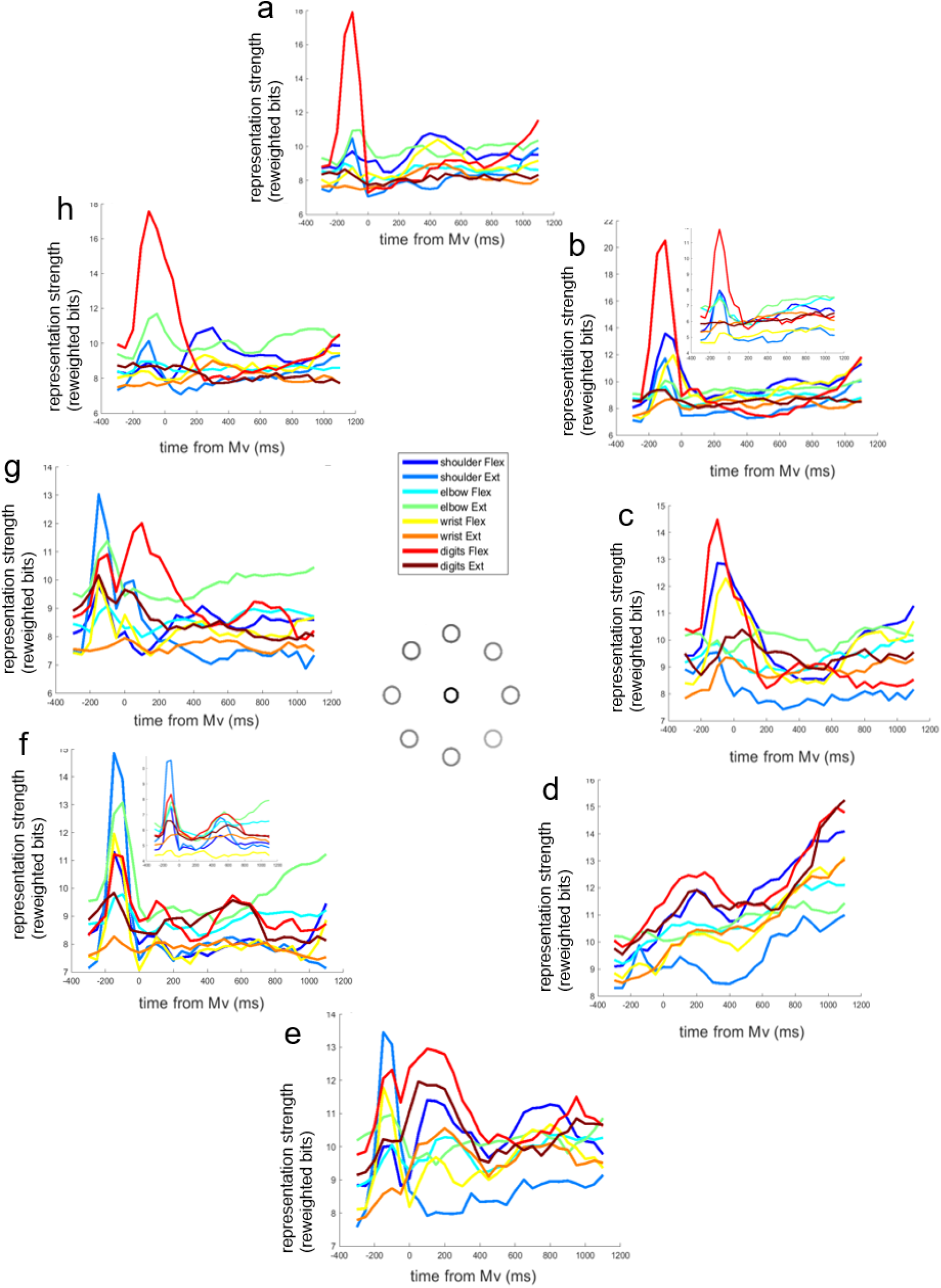
Evolution of representation strengths was movement-specific. Mutual information evolution was computed separately for eight reach directions (**a**-**h**, arranged in sequence according to actual physical reach directions). In each figure, each line was color-coded by body part and flexion/extension combinations. Insets in **b** and **f** show results from the same direction reaches from a 2-direction reach dataset obtained on a different day.

Next, we explored more thoroughly four representative diagonal directions (45º, 135º, 225º and 315º) from the 8-direction center-out dataset (Fig. 4 a-d). Common to all reach directions, more than half of the sites constantly changed their muscle representations through the movement course (blue line in the first row of Fig. 4). Only a tiny fraction of sites preserved both body part and polarity categories despite muscle change (difference between blue and purple lines in the first row of Fig. 4), while a little less than a quarter of sites preserved body part (difference between blue and red lines in the first row of Fig. 4), and roughly half of sites preserved polarity (difference between blue and yellow lines in the first row of Fig. 4). Those ratios roughly matched chance levels, indicating that the representational shifts were quite random and not particularly ‘compensatory’. For each reach direction, there was a timepoint with most representational stability, where usually 50∼60% of sites preserved the body part and flexion/extension represented (green line in the first row of Fig. 4). For most directions, this timepoint was immediately prior to movement onset, except for the 135º reach direction considered here which had a slower ramp-up. Those stability timepoints corresponded to particular layouts of the somatotopic maps (including the full map, ‘motor’ map and ‘sensory’ map, here showing only the former two in the second row of Fig. 4), which were usually vastly different for different reach directions. Maps and most represented muscle could be quite similar for similar reach directions (Fig. 4 a and d). Those most represented muscles were largely activated for those particular directions at those stability timepoints (third row in Fig. 4), though not necessarily reaching the overall EMG peak in the whole time course just yet.

**Figure 4.**
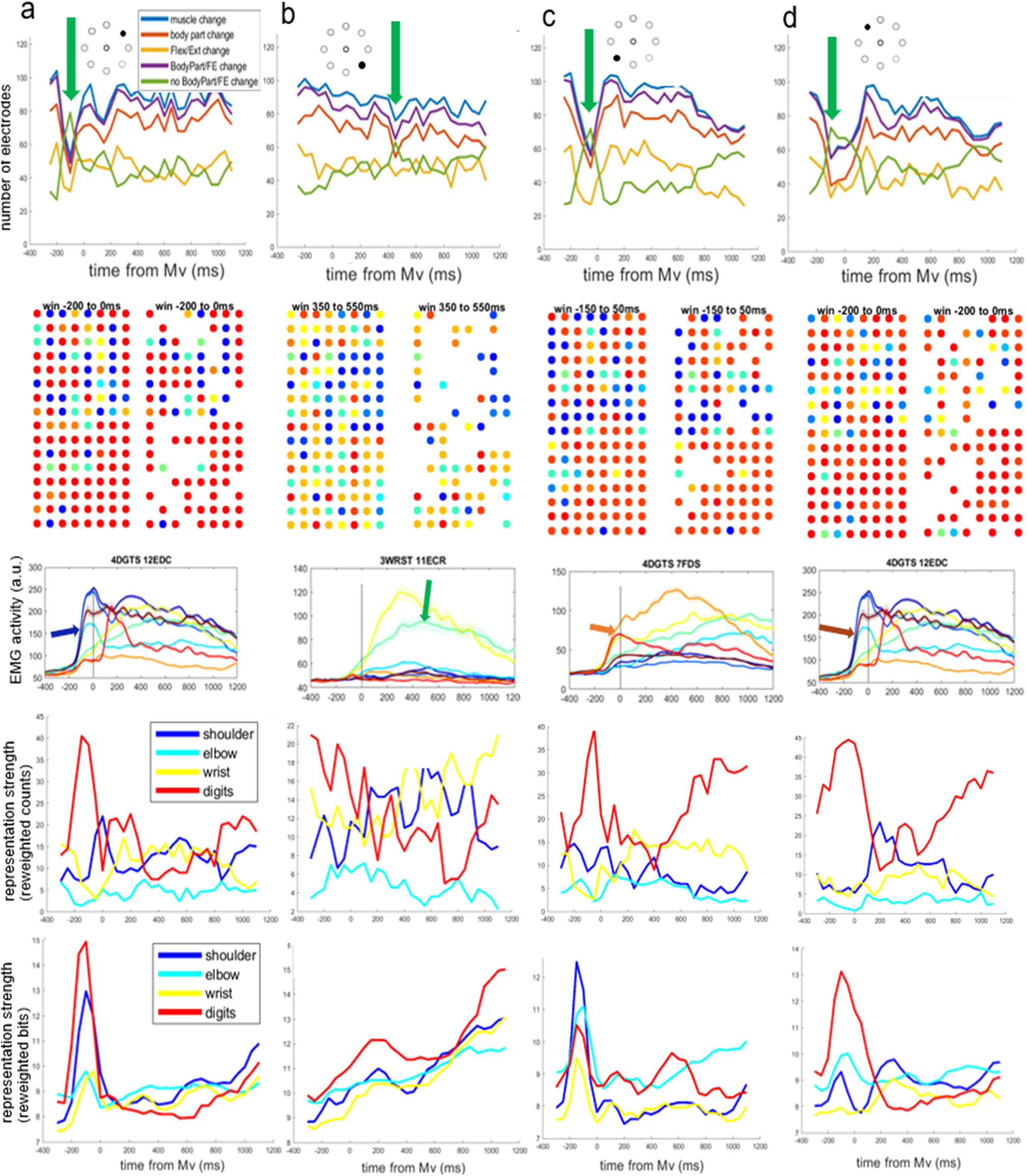
Evolution of representation (in)stability, topography and strength. Results were obtained from 8-direction center-out reaches, but only showing four diagonal directions (45º, 135º, 225º and 315º, from **a** to **d**) for clarity. Those reach directions were noted in the insets of the figures in the first row. The first row shows the instability and stability of representation from adjacent time windows. Blue lines show the number of electrodes whose most represented muscle changed in adjacent time windows; red lines show the number of electrodes whose most represented muscles in adjacent time windows were from different body parts; yellow lines show the number of electrodes whose most represented muscles in adjacent time windows were from different flexion/extension function group; purple lines show the number of electrodes whose most represented muscles changed either body part or flexion/extension; green lines show the number of electrodes whose most represented muscles kept the same body part and flexion/extension. Green arrows point to the peak of the green lines (i.e. most stability observed during the movement course). The second row shows the corresponding full somatotopic map (left) and ‘motor’ map (right) at the time point of most representational stability, color-coded using muscle colors as in the legend of Fig. 2a. The third row shows the average EMG activity for the muscle that was most represented in the full map. Each trace represented a different reach direction just as in Fig. 1, with the colored arrow pointing to the trace for the specific direction considered here. The last two rows show changes of representation strength of different body parts, computed using two different methods: the fourth row was computed in a winner-take-all sense, using adjusted electrode count as the measurement; the fifth row was computed using a MutInfo-summation method, using adjusted bits as the measurement. See Methods section for details.

Looking at the change in body part represented (last two rows of Fig. 4), again reach direction made a difference. The change of representation strength was analyzed in two ways here: in addition to the MutInfo-summation view explained above, we also counted the number of electrodes favoring each muscle (winner-take-all view). The winner-take-all (electrode count) view was in general more fluctuating than the MutInfo-summation view as expected. Interestingly, the MutInfo-summation view many times showed different trends, timing and relative ranking of body parts, compared to the winner-take-all view. For example, shoulder representation had different peaking time for 45º reach direction (Fig. 4 a, last two rows). As another example, shoulder and elbow representation was quite heavy in the MutInfo-summation view but not in the winner-take-all view, with the latter heavily dominated by digits (Fig. 4 c, last two rows). This implied that the cortical representation for shoulder and elbow were more distributed than that for digits at least during certain movements.

We also looked at data collected from another dataset of 2 reach directions (45º and 225º) on a different day, respectively in Fig. 5 a and b. The instability trends (first row in Fig. 5) and representation strength trends (last two rows in Fig. 5, especially the more stable last row) were qualitatively similar to those in Fig. 4. Somatotopic maps had varied layout, but the most represented muscle in each reach direction (in the winner-take-out sense) was the same as in Fig. 4. The consistency of results among different sessions of similar tasks indicated that the representation scheme of the same movement was relatively fixed across days.

**Figure 5.**
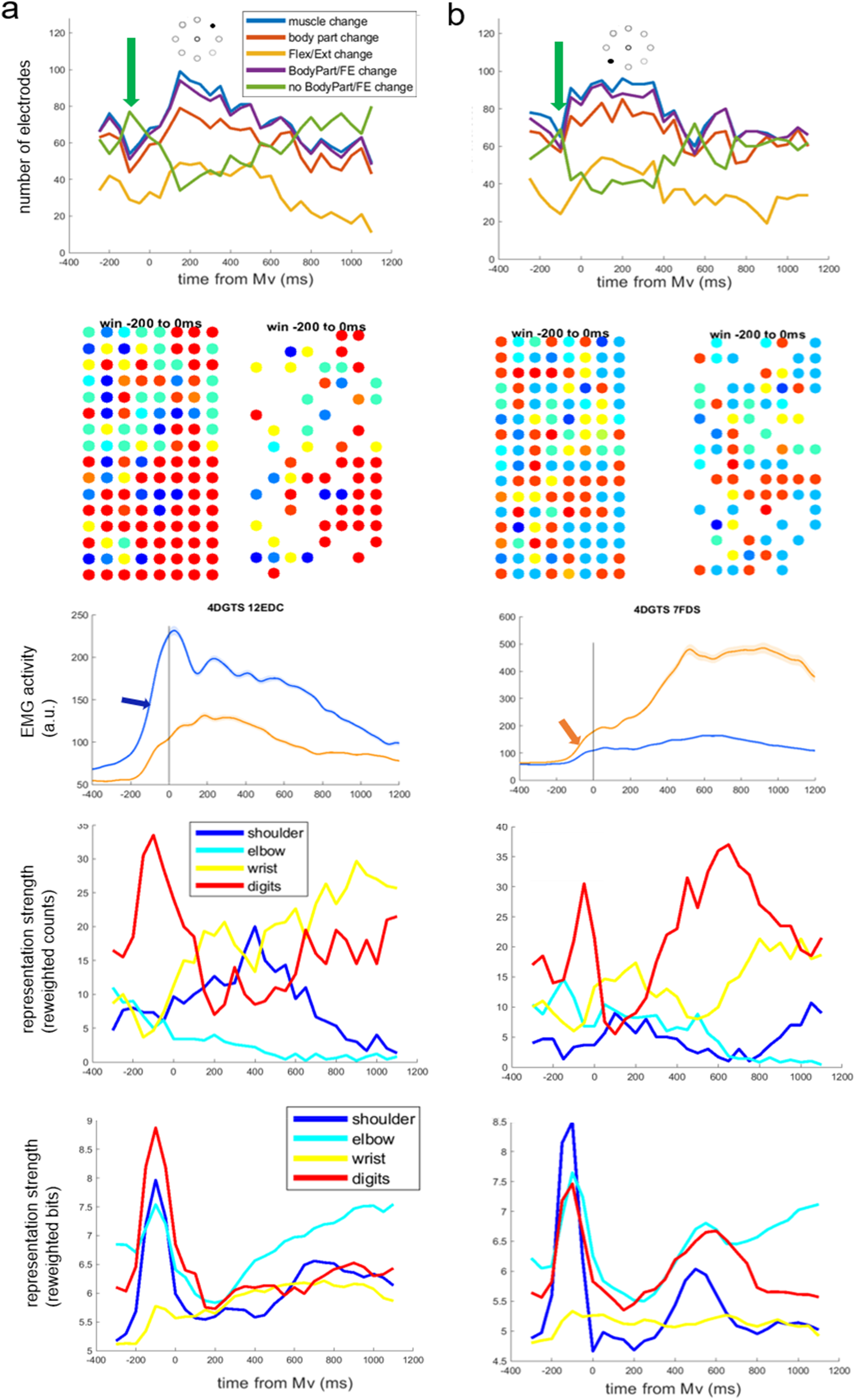
Evolution of representation (in)stability, topography and strength for another dataset. Same as Fig. 4 but obtained from another dataset of 2-direction center-out reaches (**a** for 45º, **b** for 225 º). See descriptions for Fig. 4.

### Representation for different movements could be explained by common factors

Finally, we aimed to simplify the understanding of dynamic representation for different reach directions, muscles and sites. For this purpose, we performed non-negative matrix decomposition on the tensor of mutual information (Fig. 6 a, see also methods for details), and showed that the representation for different movements could be understood using a unified set of bases. We performed the decomposition using 2∼5 factors, which produced relative errors of 0.6261, 0.6169, 0.6164, 0.6120 respectively. We chose 4 factors, considering both interpretability and faithfulness of reconstruction. Weights of the dimensions of reach direction were quite continuous for adjacent directions (Fig. 6 b). Representation for rightward and inward movements had later peaks and was slightly noisier in general (Fig. 6 c factor 2 and 1).

**Figure 6.**
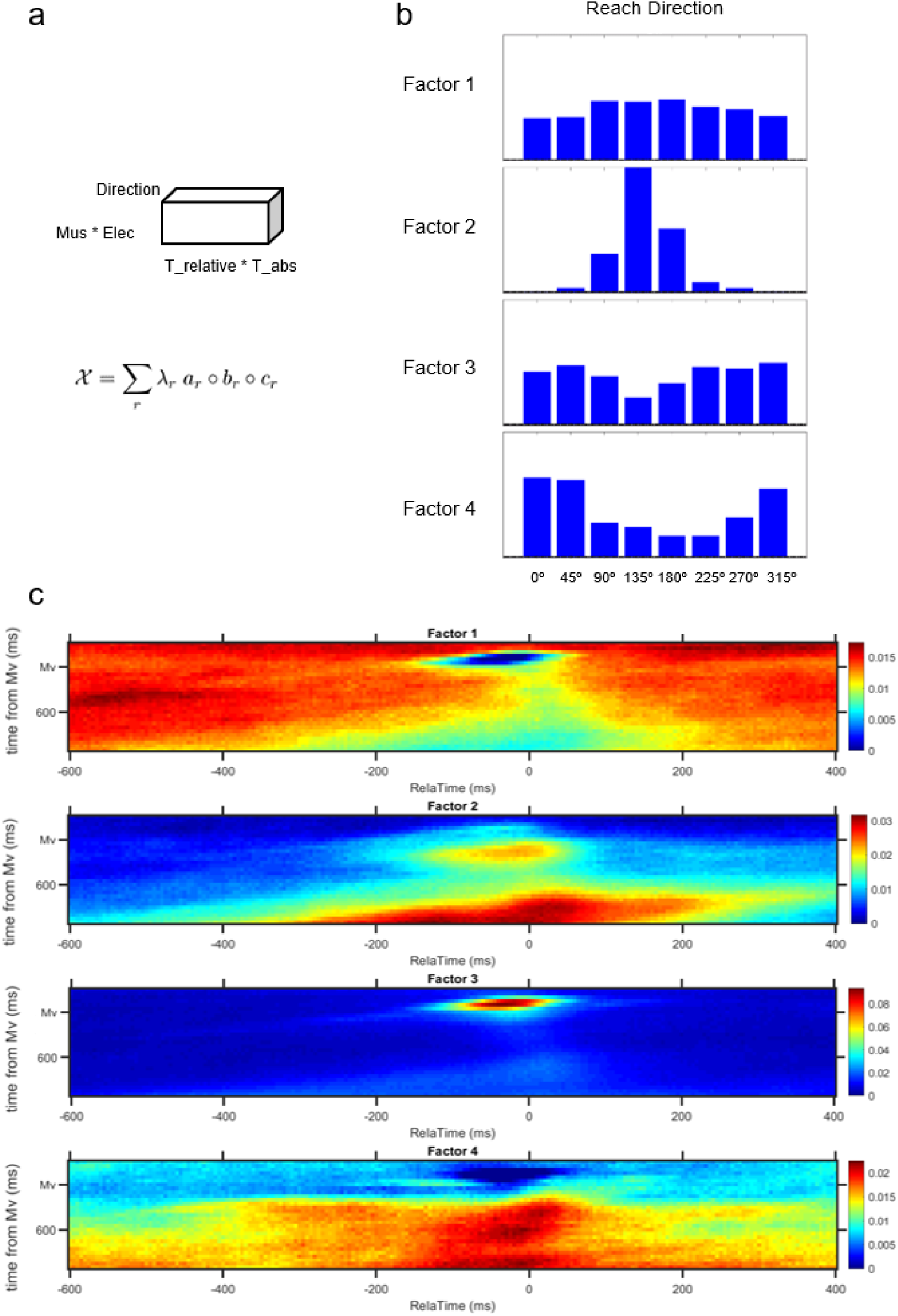
Non-negative factorization revealed common factors of representation dynamics among different reach direction. **a**. Tensor arrangement before decomposition. Three dimensions were reach direction, a merged time component, and a merged muscle & electrode component. **b**. Weights of the reach direction dimension, respectively for each of the four factors. **c**. Weights of the time dimension, respectively for each of the four factors.

Representation for most reach directions (especially outward ones) had sharper early peak which could diffuse into later movement segments (Fig. 6 c factor 3 and 4). The sharp early factor (factor 3) had an interesting spatial component as well: the lateral posterior part of the primary motor cortex was most involved in this early sharp rise of mutual information (Fig. 7 a), not only for digits muscles but also for many of the shoulder, elbow and wrist muscles. This early informative concentration later diffused to more medial and anterior parts of the cortical sheet for most muscles considered (e.g. Anterior Deltoid, Pectoralis Major, Biceps Lateral, Biceps Medial, Triceps Long head, Flexor Carpi Ulnaris, Extensor Digitorum Communis, Extensor Carpi Ulnaris), though in some cases stayed in the lateral posterior area (e.g. for Brachioradialis, Extensor Carpi Radialis and Flexor Digitorum Superficialis) (see Fig. 7 b). For those remaining rightward-inward movements which didn’t have a sharp early information rise, the slow rise of information was quite diffuse spatially, with some sparse hotspots on the cortical sheet (factor 2, spatial weights not shown here).

**Figure 7.**
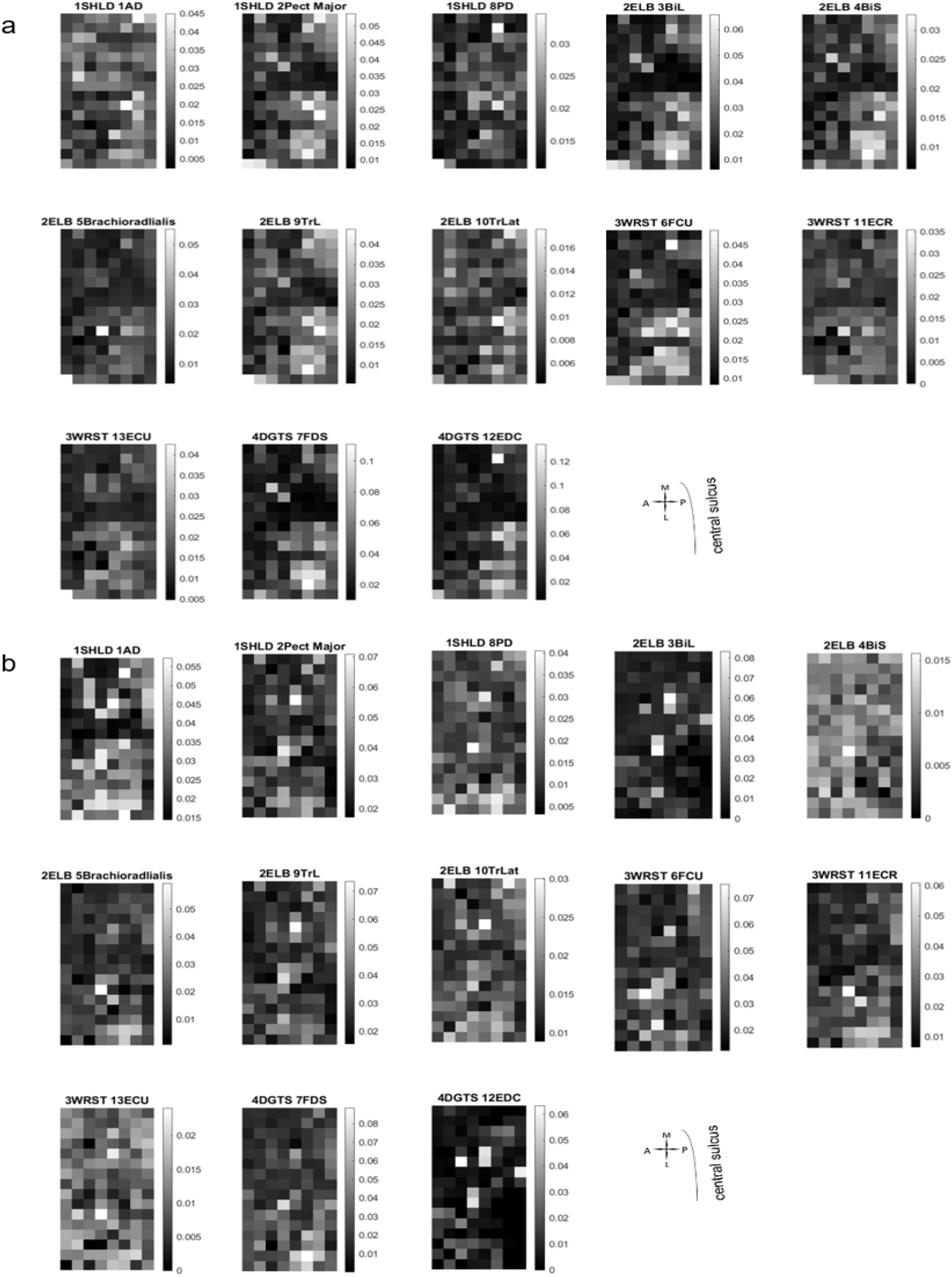
Spatial shift of information concentration. For most reach directions (except rightward-inward ones), concentration of information shifted from lateral posterior part of the cortical sheet of primary motor cortex implanted to more anterior and medial directions. **a**. Weights of the electrodes for each muscle for factor 3. **b**. Weights of the electrodes for each muscle for factor 4

## Discussion

Classic ICMS-based studies established crude somatotopic maps on the cortical sheet using short ICMS trains and, more recently, action and postural maps using long ICMS trains (Penfield & Boldrey, 1937; Woolsey, 1952; Park et al., 2001; Graziano et al., 2002), but these studies neglected a crucial temporal aspect. On the other hand, studies establishing the fluidity of movement representation generally did not consider the cortical sheet explicitly (Churchland & Shenoy, 2007; Sergio et al., 2005); in rare cases when the cortical sheet was explored, no spatial structure was found on a mesoscopic scale (Ben-Shaul et al., 2003; Hatsopoulos et al., 2007).

To our knowledge, this study is the first to discover large-scale spatial arrangements of movement representation on the cortical sheet in a time-resolved fashion. Using mutual information as the measurement of representation strength, we confirmed the ever-evolving nature of movement representation (more specifically, muscle representation) and illustrated each representation snapshot on the cortical sheet. Despite the distributed and overlapping nature of muscle encoding on the cortical sheet, a clear distal-proximal gradient was present throughout movement course. Furthermore, these dynamic representations on the cortex were movement-specific, with varying times of stability and topographic maps, possibly serving functional needs.

### Temporal representation

Temporally, we characterized a special timepoint in the course of representational evolution — movement onset where the representations for individual muscles were usually strongest and most stable immediately prior to movement onset. This unique timepoint echoes to what others have found. Sergio et al. (2005) discovered that with the center-out movement task, single units developed directional tuning upon force onset, which then continuously changed directional tuning. Using a similar task, Ben-shaul et al (2003) also discovered changing tunings of single units, which was particularly different between holding and execution. Those affirmative results were encouraging especially considering the complementary nature of their formulation of tuning versus our formation of representation (which will be discussed in more detail in later part of Discussion).

Another prominent feature for the temporal evolution of representation was its movement-specificity. We found that different movement directions had different representational changes; in other words, specific recruitment patterns of muscles strongly covaried with its instantaneous cortical representation. Moreover, the stabilized maps had strong representations for muscles that were heavily recruited at those times for those particular reach directions.

So what could be the underlying reason for representations to be movement-specific? Larger mutual information was not a trivial by-product from simply having larger EMG activity, since (1) in our observation, the largest EMG activity didn’t always coincide in time exactly with the largest mutual information; and (2) in theory, mutual information is reparameterization-independent (Kraskov et al., 2004), so simple scaling of the amplitudes of the muscle signals and/or neural signals would not change their mutual information. Instead, this reorganized representation likely reflected actual *functional needs* – the need to represent finely the muscles that were mostly needed at this particular timepoint. This explains why we saw non-canonical time courses for representational stability in direction 135º (Fig. 4 b first row), where representation was stabilized as late as 450ms; most likely it was due to the need for later muscle activation (e.g. Extensor Carpi Radialis). There exists additional evidence in support for the argument for functional needs. Sergio et al. (2005) explored another task in the same study mentioned above, which was isometric (requiring monotonic force ramping thus exhibiting monotonic muscle activity). Curiously, no change in tunings was observed once tunings were established upon force onset. Perhaps these relatively stable tunings in this task of theirs were precisely due to the scarce need of functional change of representation in this scenario. In contrast, the center-out movement task in their and our studies, where muscles tended to exhibit tri-phasic activities (Forget & Lamarre, 1987), required more stages of varying representation in order to fulfill varying recruitment needs. It would be illuminating to conduct our analyses with their isometric task to confirm this hypothesis.

### Spatial representation

Regarding space, while Hatsopoulos et al. (2007) and Ben-shaul et al. (2003) discovered that the same site or neighboring sites tended to have relatively similar directional tuning, there was no systematic organization over cortical sheet on the mesoscopic scale. This study, in contrast, discovered that muscle representations were very clustered on a scale of tens of µms, also conforming to the proximal-distal (body part) gradient. This stark disparity was likely mostly due to the different choices of tuning features (preferred direction in the kinematic view versus muscle in the kinetic view). Despite constant fluidity, muscle representation is more spatially-organized on the cortical sheet than preferred directions, possibly because the former was to some extent developmentally hard-wired following gradients due to different timings of circuit development (in spinal cord and brain stem) for proximal versus distal limbs (Martin et al., 2005), while the latter was predominantly secondarily formed based off interactions with the physical surroundings, thus having to evolve under pre-existent representation constraints (Aflalo & Graziano, 2006).

As a side note, in a spatially-continuous manner, we discovered interleaved ‘motor’, ‘sensory’ and ‘simultaneous’ locations for the same muscles (Fig. 2). This perhaps seems puzzling at first glance, considering all recorded sites were in primary motor cortex, but is not too surprising in the literature (for a review, see Hatsopoulos & Suminski, 2011). Potentially, those ‘simultaneous’ and ‘sensory’ locations could serve in predictive control (Flament & Hore, 1988), feedback control (Pruszynski et al., 2011), kinesthetic perceptions (Naito et al., 2011) and movement observation (Suminski et al., 2009). The spatial continuity in those different lag groups pointed to the integrative and collaborative nature of motor representation.

### Interplay between spatial and temporal representation

Not only was muscle representation spatially organized on the cortical sheet throughout movement, there existed interplay between cortical space and time in movement. Firstly, we discovered enlarged proximal-distal gradient encompassing more diverse body parts upon movement onset (Fig. 2). Interestingly, Ben-shaul et al. (2003) found that neighboring electrodes exhibited bigger tuning disparity during movement compared to preparation (75° vs. 45° of tuning disparity during movement vs. during preparation), which corresponded to an increase in representation diversity upon movement onset in our study. The preservation of proximal-distal gradient could potentially help to maintain a relatively stable representation on the mesoscopic spatial scale, thus reducing the efforts of having to make drastic movement-dependent changes in various anatomical and functional connections.

Secondly, we found that close to movement onset, the most informative locations on the cortical sheet were at the lateral-posterior part of the primary motor cortical sheet probed, for many muscles distributed across forelimb; this information concentration later diffused to other locations on the sheet. To probe for those patterns, we intentionally kept muscle and electrode together in one dimension in the non-negative tensor decomposition, which allowed patterns between muscles and electrodes to form naturally. As commonality of spatial component among muscles was not forced in the dimensionality reduction, it is more meaningful that we discovered similar spatial components for different muscles, including distal and proximal muscles alike. This early ‘universal’ information hotspot in primary motor cortex we discovered here might indicate that at the early stage of movement this area was most direct in dictating muscle activities. For example, we know that there were more cortico-motoneuronal (CM) cells in caudal primary motor cortex, which made monosynaptic connections to motoneurons in the ventral horn; in comparison, the rostral area had fewer CM cells but more corticospinal (CST) neurons, mostly connecting to interneurons in the spinal cord (Rathelot & Strick, 2009).

Importantly, the concentration of CM cells in caudal primary motor cortex not only connected to motoneurons controlling distal muscles, but also those controlling proximal muscles (Rathelot & Strick, 2009), which was reminiscent of what we found here. While further studies will be needed to elucidate the origin of this phenomenon, there are definitely advantages to such an arrangement. An early concentrated information hub could facilitate early coordination of muscle synergies, as different combinations of muscle recruitments can be flexibly used to achieve the same limb movement.

Lastly, an interesting observation is that the relative representation strength in the winner-take-all view and summation view did not always agree, especially as time evolved (Fig. 4 and 5, last two rows). This fact implied that information was distributed to different extents for different body groups, and for different timepoints throughout movement. Representation for digits tended to be the more concentrated in single electrodes than for elbow or shoulder, perhaps due to less coordinative requirements for distal joints compared to connective joints. This observation is to some extent in line with the classical horseshoe shape of upper limb representation, where larger distal and intermediate representations encircled smaller distal representation (Park et al., 2001).

### Measurement of representation

Crucial differences between our study and previous studies lie in our choice of representational measurement. Representation, or mapping, is supposed to be a relationship between (at least) two variables, which should be computed (at least) at 2nd-order. Traditionally, people used (stronger) neural activation to serve as proxy of (stronger) representation, studied at each movement or effector group, which could result in fragmented or biased understanding (e.g. neglecting inhibitory effects when studying tuning). In contrast, we directly measured the inter-dependence between effector of interest and neural activation, which is a 2nd-order measurement that directly speaks to the representation issue at hand. Furthermore, we chose to use mutual information in the continuous formulation as the measurement of inter-dependence (Kraskov et al., 2004; Holmes & Nemenman, 2019), which offers comprehensive characterization of representation encompassing all activity levels of the two variables without arbitrary binning or grouping. Despite popularity in other quantitative fields, this method of continuous mutual information calculation had just started to be utilized in neuroscience (Srivastava et al., 2017). We think the advantages of this method outweigh its computational encumbrance at least in studies of strongly continuous signals like ours, and encourage further attempts in the neuroscience community. Lastly, there was a potential caveat in our manipulation of mutual information after it was computed — to calculate the total representation in the electrodes studied, we summed up the mutual information bits. Summation of mutual information here was not entirely justified, as we could not guarantee that the information obtained from different electrodes were not redundant. However, as it is hardly tractable to compute mutual information between high-dimensional variables, we hoped to provide a simple estimate for the total information in the system, assuming the level of non-redundancy did not differ too much among muscles or among electrodes. It would be useful to re-evaluate those hypotheses once techniques mature.

To summarize, we demonstrated the multiplexed and distributed nature of muscle representation on the cortical sheet of primary motor cortex, from the perspective of spatio-temporal information evolution. Representation constantly evolved in the course of single reaches and among different movements, but was mostly stable upon movement onset, likely reflecting functional needs. We also discovered temporal evolutions in the spatial gradient of body parts and in the information hotspot, emphasizing the added value of studying simultaneously space on cortical sheet and temporal evolution. Future directions include characterizing the representation stability at different spatial scales, linking evolving local networks to evolving representation at single sites, and developing online detection methods for informative sites to improve brain-machine-interfaces.

## Acknowledgments

We thank Caroline M. Holmes and Ilya Nemenman for helpful discussions regarding the modified KSG mutual information estimator. We thank Stephanie Palmer, Jason MacLean and Yali Amit for helpful suggestions regarding analyses. We also thank the University of Chicago research computing center for providing computing resources for intensive computations of mutual information. This study is supported by NIH R01 NS111982.

